# m^7^G cap-eIF4E interaction stimulates polysome formation by enhancing first-round initiation kinetics

**DOI:** 10.1101/2021.12.29.474420

**Authors:** Lexi Sun, Anthony Gaba, Hongyun Wang, Xiaohui Qu

## Abstract

Translation in eukaryotic cells occurs predominantly through a 7-methylguanosine (m^7^G) cap-dependent mechanism. m^7^G cap interactions with eukaryotic initiation factor 4E (eIF4E) facilitates 43S recruitment to the mRNA 5’ end and enhances the translation efficiency of mRNA. However, it remains poorly understood how m^7^G cap-eIF4E interactions affect polysome formation kinetics. Here, we examine the role of the m^7^G cap in polysome formation by utilizing a single-molecule approach to track individual ribosomes during active translation. Translation was monitored in wheat germ extract with capped and uncapped synthetic mRNAs and in HeLa extract with purified human eIF4E titration. The presence of the m^7^G cap and the supplementation of eIF4E to eIF4E-deficient extract enhanced the kinetics of the first initiation event of polysomes. Subsequent to the first initiation event, efficient polysome-forming initiation events occurred independent of mRNA m^7^G capping status and eIF4E concentration. Our results indicate that m^7^G cap-eIF4E interactions in wheat germ and HeLa extracts promote polysome formation by enhancing first-round initiation kinetics. The dynamics of individual translation events on polysomal mRNAs suggest that first-round initiation events activate mRNAs for efficient subsequent rounds of polysome-forming initiation.

## Introduction

The cap-dependent translation initiation pathway in eukaryotic cells occurs through a scanning mechanism that begins with association of a 40S ribosomal subunit with a ternary complex (eukaryotic initiation factor (eIF) 2, GTP, and methionyl initiator tRNA) and the eIFs 1, 1A, 3, and 5, which collectively form a 43S pre-initiation complex (1). The 5’-terminal 7-methylguanosine (m^7^G) cap structure facilitates 43S recruitment to the mRNA 5’ end, from which the 43S scans the mRNA 5’-untranslated region (UTR) in a 5’ to 3’ direction (2). 43S recognition of a start codon stops the scanning 43S and leads to release of eIFs, PIC structural rearrangements, and 60S subunit joining to form the 80S ribosome (1, 2), which then proceeds to the peptide elongation stage. Elongation continues until the ribosome reaches a termination codon and incorporates a release factor which releases the ribosome from the mRNA (3). The m^7^G cap functions through interactions with the cap binding protein eIF4E, which together with eIF4A and eIF4G forms the heterotrimeric complex, eIF4F. Interactions with eIF4F, the poly (A) binding protein (PABP), and mRNA are thought to mediate an mRNA closed loop structure that activates mRNAs for efficient translation (4, 5). The translation dynamics mediated by these interactions remain incompletely understood.

If initiation events occur when a previously initiated ribosome is actively engaged in elongation, more than one ribosome can simultaneously translate the same mRNA and form a polysome (6-9). Polysome formation can be detected by polysome profiling using sucrose gradient fractionation (10), ribosome profiling (11), cryoEM (12, 13), and *in vivo* fluorescence imaging (14, 15). However, polysome formation dynamics remain poorly understood because the resolution of these techniques limits their ability to dissect the kinetics of individual ribosomes that form polysomes. To access the dynamics of polysome formation, there is a need for new methods that can resolve the kinetics of individual translating ribosomes on single mRNA molecules.

We previously developed a single-molecule assay that enables tracking of single ribosome translation kinetics during active cap-dependent translation (16-18). The assay is based on detection of fluorescently-labeled antibody binding to nascent N-terminal-tagged polypeptide during cell-free translation. The versatility of the assay allows its application to diverse extract-based cell-free translation systems, including wheat germ, yeast, and rabbit reticulocyte systems (18). Here by utilizing this single-molecule assay with wheat germ and HeLa extracts, we tracked the kinetics of individual translating ribosomes during polysome formation in plant and mammalian translation systems. Our single-molecule analyses uncover a first-round initiation-specific role for m^7^G cap-eIF4E interactions that activates mRNAs for efficient polysome formation.

## Results

### Single-molecule detection and quantification of polysome formation kinetics

Our previously established *in vitro* single-molecule assay enables tracking of individual translation events by single ribosomes during active cap-dependent translation (17, 18). The assay utilizes cell-free translation and is compatible with diverse systems, including cell extracts from yeast, wheat germ, and rabbit reticulocyte lysates (18). To apply the assay to investigate polysome formation kinetics, we used cell-free translation with wheat germ extract (WGE) and synthetic m^7^G-capped and polyadenylated mRNA (*m*^*7*^*G-3xFLAG-Fluc-pA*) that encodes an N-terminal 3xFLAG tagged firefly luciferase (Figure 1A). 3’ end biotinylation of the *m*^*7*^*G-3xFLAG-Fluc-pA* mRNA enables mRNA tethering to a streptavidin-coated detection surface via biotin-streptavidin interaction in a flow channel (17). By flushing into the flow channel translation mix containing WGE supplemented with 134 nM Cy3 labeled anti-FLAG antibody (Cy3-αFLAG), translation of 3’ end-tethered *m*^*7*^*G-3xFLAG-Fluc-pA* mRNA exposes the nascent FLAG-tagged luciferase peptide (Fluc_FLAG_) from the ribosome tunnel and allows Cy3-αFLAG interaction with the FLAG tag (Figure 1B).

**Figure 1.**
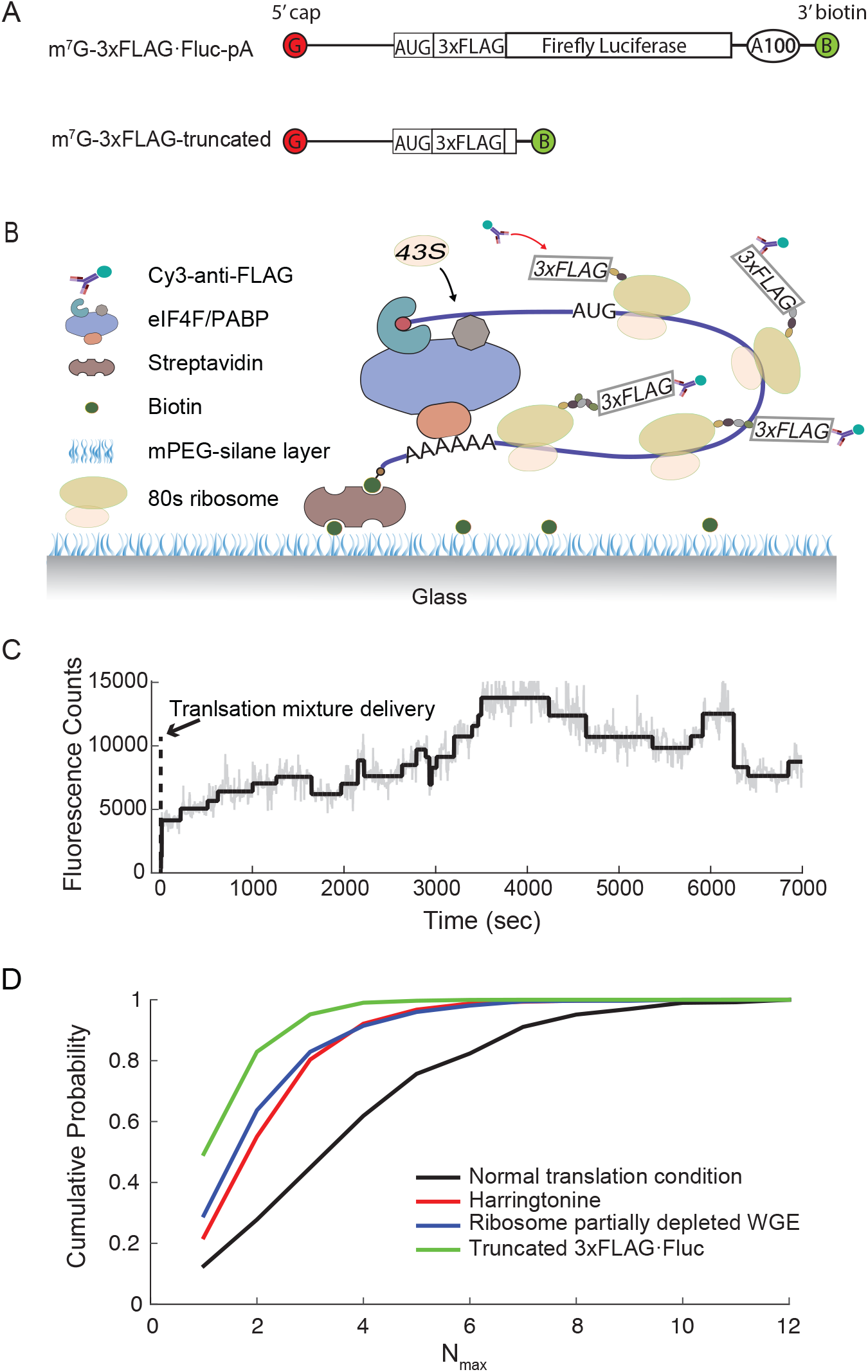
A reconstituted cell-free translation system for single-molecule analysis of polysome formation kinetics. (*A*) Schematic representation of full length synthetic m^7^G-capped and polyadenylated mRNA that encodes an N-terminal 3xFLAG tagged firefly luciferase (top), and a truncated mRNA with shortened firefly luciferase coding region (bottom). (*B*) Schematic of the single-molecule assay, which allows real-time detection of Cy3-αFLAG binding to nascent 3xFLAG generated during active *in vitro* translation of 3’ end surface anchored reporter mRNA. (*C*) A representative single molecule trajectory for WGE translation of an individual *m*^*7*^*G-3xFLAG-Fluc-pA* mRNA. The early increase, indicated by arrow and denoted as time 0, results from the delivery of cell-free translation mix supplemented with Cy3-αFLAG. Raw and digitized data are shown in gray and black, respectively. (*D*) Cumulative probability distribution of maximum number of co-existing translating ribosomes on single mRNAs under four different translation conditions: i) the normal condition (WGE translation of *m*^*7*^*G-3xFLAG-Fluc-pA* mRNA), ii) with addition of harringtonine, iii) translation using ribosome partially depleted WGE, and iv) translation of truncated *m*^*7*^*G-3xFLAG-truncated* mRNA.

Under the above experimental condition, Cy3-αFLAG binding to nascent Fluc_FLAG_ occurs rapidly with a time constant of less than 4 sec, is highly specific with a specific to non-specific binding ratio greater than 20-fold, and is stable for at least 30 min. (18). To further confirm Cy3-αFLAG specific binding to nascent 3xFLAG-Fluc in WGE, either 1mM puromycin or vehicle control was flushed into channels after 5 min of *m*^*7*^*G-3xFLAG-Fluc-pA* mRNA translation in WGE containing Cy3-αFLAG. Puromycin accelerated the rate of Cy3-αFLAG disappearance by 12-fold (Figure S1). The faster rate of Cy3-αFLAG dissociation with puromycin is consistent with the known ability of puromycin to enter the peptidyl transferase center A site and form a peptide-puromycin product that terminates translation prematurely and dissociates translating ribosomes from mRNA (19-21). Taken together, the results indicate that individual Cy3-αFLAG binding and dissociation events serve as faithful trackers of initiation, elongation, and termination of individual translating ribosomes.

Cy3-αFLAG interactions with the N-terminal 3xFLAG tag on individual nascent Fluc_FLAG_ peptide were analyzed as trajectories showing Cy3 fluorescence changes for individual mRNA molecules (Figure 1C). Each instantaneous fluorescence increase or decrease step represents the onset and termination of peptide elongation by individual ribosomes, respectively. The trajectories show Cy3-αFLAG binding events with accumulating clusters of Cy3-αFLAG-Fluc_FLAG_ complexes on individual *m*^*7*^*G-3xFLAG-Fluc-pA* mRNAs, suggesting that multiple ribosomes simultaneously translated individual mRNAs and formed polysomes. To further examine Cy3-αFLAG binding events for polysome formation, we tested for translation-dependent clusters of Cy3-αFLAG-Fluc_FLAG_ complexes by diminishing translation activity with (i) WGE containing partially depleted ribosomes, (ii) translation mix supplemented with the initiation inhibitor harringtonine, and (iii) a truncated *m*^*7*^*G-3xFLAG-Fluc-pA* mRNA with a coding region shortened to 259 nt (*m*^*7*^*G-3xFLAG-truncated*) (Figure 1A). To compare the extent of polysome formation between the different translation conditions, the maximum numbers of simultaneously bound Cy3-αFLAG antibodies on single mRNA molecules (Nmax) were measured (Figure 1D). Compared to the control translation condition, each condition with diminished translation activity showed cumulative probability distributions with reduced Nmax values. These results support the interpretation that the accumulated Cy3-αFLAG binding events in trajectories (Figure 1C) represent polysomes and can be used to monitor the dynamics of polysome formation.

### m^7^G cap-eIF4E interactions specifically stimulate the first round of initiation

Distinct stages of the canonical m^7^G cap-dependent initiation pathway have been characterized (22). However, effects of these individual steps on polysome formation dynamics remain poorly understood. To investigate the influence of specific initiation steps on polysome formation, Cy3-αFLAG binding events were analyzed in translation reactions that were defective for distinct initiation steps. Specifically, effects of m^7^G cap-eIF4E and poly(A) tail-PABP interactions were tested with *3xFLAG-Fluc*-encoding mRNAs that either contained or lacked an m^7^G cap and poly(A) tail (*m*^*7*^*G-3xFLAG-Fluc-pA, m*^*7*^*G-3xFLAG-Fluc, ppp-3xFLAG-Fluc-pA*, ppp*-3xFLAG-Fluc*) (Figure 2A). Effects of 43S scanning were tested by impeding scanning with a 14 base pair 100% GC-rich hairpin (ΔG = -35.30 kcal/mol) positioned 90 nt from the mRNA 5’ end and 87 nt from the *3xFLAG-Fluc* AUG start codon (*m*^*7*^*G-hp-3xFLAG-Fluc-pA*) (Figure 2A).

**Figure 2.**
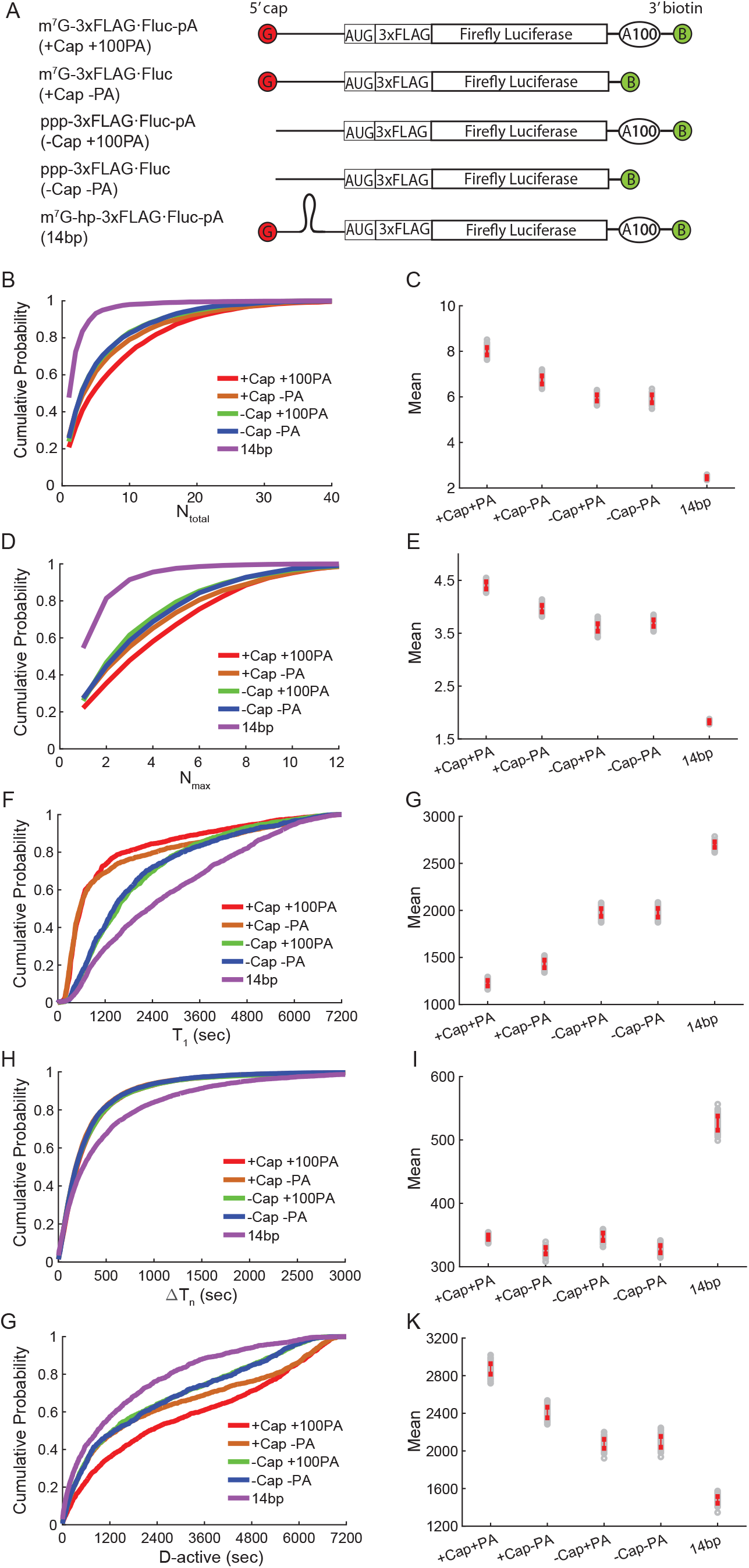
Effects of 5’ m^7^G cap, 3’ poly(A) tail, and mid-5’ UTR stable hairpin on polysome formation in WGE translation of *3xFLAG-Fluc* mRNA. (*A*) Constructs of *3xFLAG-Fluc* mRNA variants that contain or lack 5’ m^7^G cap, 3’ poly(A) tail, and a mid-5’ UTR stable hairpin. (*B, C*) Cumulative probability distribution (*B*) and mean values (*C*) of the total number of Cy3-αFLAG binding per trajectory (Ntotal). Mean ± S.E.: 8.0 ± 0.2 (+Cap+100PA), 6.7 ± 0.2 (+Cap-PA), 6.0 ± 0.1 (-Cap+100PA), 5.9 ± 0.2 (-Cap-PA), 2.45 ± 0.05 (14bp). Numbers of trajectories analyzed: n = 2192 (+Cap+100PA), 2060 (+Cap-PA), 1865 (-Cap+100PA), 1381 (-Cap-PA), and 3405 (14bp). (*D, E*) Cumulative probability distribution (*D*) and mean values (*E*) of the maximum number of simultaneously bound antibodies (Nmax). Mean + S.E.: 4.41 ± 0.07 (+Cap+100PA), 3.97± 0.06 (+Cap-PA), 3.61 ± 0.07 (-Cap+100PA), 3.69 ± 0.06 (-Cap-PA), 1.83 ± 0.02 (14bp). Numbers of trajectories analyzed are the same as in (*B, C*). (*F, G*) Cumulative probability distribution (*F*) and mean values (*G*) of the duration from translation mix delivery to the first Cy3-αFLAG binding event (T1). Mean ± S.E. (sec): 1230 ± 30 (+Cap+100PA), 1430 ± 40 (+Cap-PA), 1980 ± 40 (-Cap+100PA), 1980 ± 50 (-Cap-PA), 2700 ± 30 (14bp). Numbers of trajectories analyzed are the same as in (*B, C*). (*H, I*) Cumulative probability distribution (*H*) and mean values (*I*) of time lags between the onsets of Cy3-αFLAG binding events in clusters of translation activity (ΔTn). Mean + S.E. (sec): 346 ± 3 (+Cap+100PA), 325 ± 5 (+Cap-PA), 347 ± 6 (-Cap+100PA), 328 ± 6 (-Cap-PA), 530 ± 10 (14bp). Numbers of events analyzed: n = 15007 (+Cap+100PA), 11560 (+Cap-PA), 8895 (-Cap+100PA), 6618 (-Cap-PA), and 4173 (14bp). (*J, K*) Cumulative probability distribution (*J*) and mean values (*K*) of the duration from the first Cy3-αFLAG binding event to the last Cy3-αFLAG binding event per cluster of translation activity (D-active). Mean ± S.E. (sec): 2870 ± 60 (+Cap+100PA), 2410 ± 60 (+Cap-PA), 2080 ± 50 (-Cap+100PA), 2100 ± 60 (-Cap-PA), and 1480 ± 40 (14bp). Numbers of events analyzed: n = 1821 (+Cap+100PA), 1565 (+Cap-PA), 1487 (-Cap+100PA), 1039 (-Cap-PA), and 1610 (14bp).

Trajectories (Figure S2A) showed that the m^7^G cap and hairpin structure each altered Cy3-αFLAG binding trajectories by regulating either the onset time or the number of Cy3-αFLAG binding events per trajectory. The effect of the poly(A) tail on trajectories were less significant than other conditions. To assess polysome size at a given timepoint, we quantified the mean numbers of simultaneously bound Cy3-αFLAG on single mRNA molecules (Figure S2B). The time-resolved kinetics of Cy3-αFLAG binding indicated that the m^7^G cap and poly(A) tail each increased polysome formation kinetics by different extent, whereas the 5’ UTR hairpin reduced polysome formation kinetics (Figure S2B). These observations suggest that the m^7^G cap, poly(A) tail, and 5’ UTR hairpin each affected the dynamics of polysome formation through distinct mechanisms.

To dissect the mechanism of polysome formation on single mRNA, we quantify the single-molecule trajectories from five different perspectives (Figure S3): i) the total number of Cy3-αFLAG binding (Ntotal), a measure of total translation activity per mRNA; ii) the maximum number of simultaneously bound antibodies (Nmax), a measure of the extent of polysome formation; iii) the duration from translation mix delivery to the first Cy3-αFLAG binding event (T1), a measure of how fast each mRNA is activated for protein synthesis; iv) time lags between the onsets of Cy3-αFLAG binding events in clusters of translation activity (ΔTn), a measure of ribosome recruitment kinetics during polysome formation; and v) the duration from the first Cy3-αFLAG binding event to the last Cy3-αFLAG binding event per cluster of translation activity (D-active), a measure of how long an mRNA is engaged in the polysome formation phase.

The data (Figures 2B-E) show that the m^7^G cap and poly(A) tail increased Ntotal and Nmax, indicating that on individual mRNAs, m^7^G cap-eIF4E interactions and poly(A) tail-PABP interactions promoted initiation and polysome formation. Analysis of T1 showed that the m^7^G cap and poly(A) tail decreased the Cy3-αFLAG first arrival time (Figures 2F and 2G). Interestingly, analysis of ΔTn showed that neither the m^7^G cap nor poly(A) tail changed the time lags between the onsets of subsequent Cy3-αFLAG binding events (Figures 2H and 2I). This unexpected and stark contrast of the strong dependence of T1 vs. the independence of ΔTn on the presence of m^7^G cap and poly(A) tail indicates that the mRNA activation process and the active polysome formation phase are regulated by different mechanisms. Specifically, cap-eIF4E and poly(A) tail-PABP interactions increased the *3xFLAG-Fluc* mRNA translation efficiency predominantly by expediting the mRNA activation process without detectable impact on ribosome recruitment dynamics during the polysome formation phase. Analysis of D-active showed that the duration of polysome formation phase was prolonged in the presence of m^7^G cap and poly(A) tail, indicating that m^7^G cap-eIF4E interactions and poly(A) tail-PABP interactions helped to maintain the mRNA in the polysome formation phase (Figures 2J and 2K). In addition, the ability of the poly(A) tail to increase Ntotal, Nmax, T1, and D-active required the presence of the m^7^G cap (Figures 2B-G, 2J, and 2K), indicating that poly(A) tail-mediated regulation was m^7^G cap-dependent. In contrast, *m*^*7*^*G-hp-3xFLAG-Fluc-pA* showed reduced initiation and polysome formation in all five kinetic analyses, indicating that the 5’ UTR hairpin inhibited ribosome scanning to the *3xFLAG-Fluc* start codon in every single round of translation.

Altogether, the analysis of Ntotal, Nmax, T1, ΔTn and D-active revealed several important insights of the *3xFLAG-Fluc* mRNA translation process: i) the mRNA translation can be divided into inactive and active ribosome recruitment phase; ii) m^7^G cap-eIF4E interactions and poly(A) tail-PABP interactions stimulated initiation and polysome formation both by expediting the transition of mRNA from the inactive to active ribosome recruitment state and by maintaining the active phase; iii) the cap-eIF4E interaction was the main regulator of this process and poly(A) tail-mediated regulation was m^7^G cap-dependent; and iv) ribosome recruitment dynamics during the active phase is independent of both m^7^G cap-eIF4E and poly(A) tail-PABP interactions.

### Human eIF4E specifically stimulates first-round initiation events in HeLa cell extract

To examine effects of m^7^G cap-eIF4E interactions on mammalian translation dynamics, HeLa cell extract was applied to the *in vitro* single-molecule assay and was supplemented with either 200 nM or 500 nM purified human eIF4E. Compared to translation reactions without eIF4E supplementation, 200 nM and 500 nM eIF4E supplementation increased Ntotal by 17% and 42%, respectively (Figures 3A and 3B). These observations indicate that *m*^*7*^*G-3xFLAG-Fluc-pA* mRNA translation in this HeLa extract was limited by eIF4E availability. To assess effects of eIF4E supplementation on polysome dynamics in HeLa extract, Nmax were measured and showed that 200 nM and 500 nM eIF4E supplementation increased Nmax by 8% and 30%, respectively (Figures 3C and 3D). 500 nM eIF4E supplementation to translation reactions with uncapped *ppp-3xFLAG-Fluc-pA* mRNA showed Ntotal and Nmax values that were similar to those with capped *m*^*7*^*G-3xFLAG-Fluc-pA* mRNA in the absence of eIF4E supplementation (Figures 3A-D). Collectively, these results indicate that in this HeLa extract, m^7^G cap-eIF4E interactions stimulate *m*^*7*^*G-3xFLAG-Fluc-pA* mRNA translation and polysome formation.

**Figure 3.**
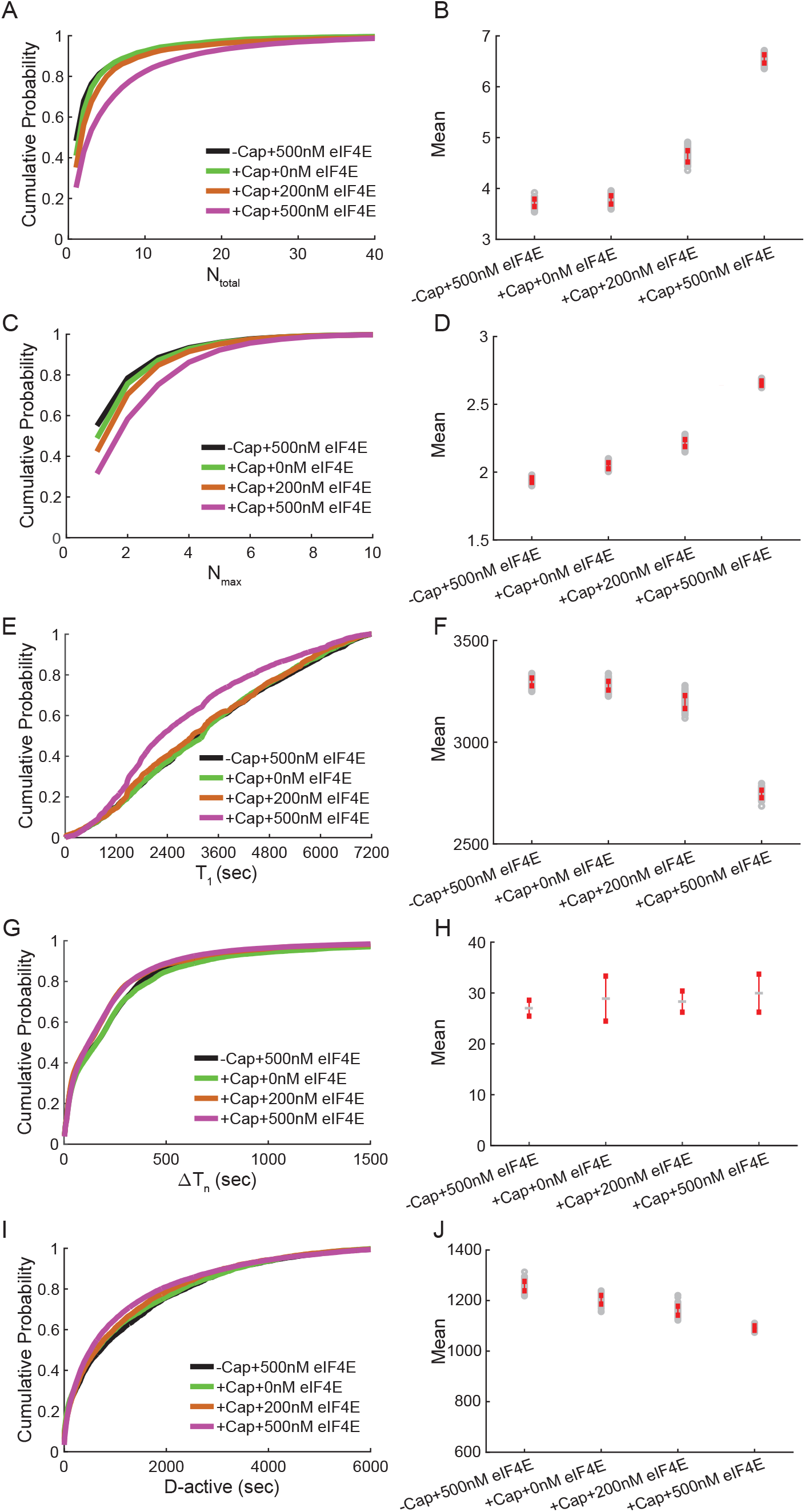
Effect of eIF4E on polysome formation in HeLa extract translation of *3xFLAG-Fluc* mRNA. (*A, B*) Cumulative probability distribution (*A*) and mean values (*B*) of Ntotal. Mean ± S.E.: 3.72 ±0.07 (-Cap+500nM eIF4E), 3.78 ± 0.08 (+Cap+0nM eIF4E), 4.6 ± 0.1 (+Cap+200nM eIF4E),6.55 ± 0.08 (+Cap+500nM eIF4E). Numbers of trajectories analyzed: n = 8329 (-Cap+500nM eIF4E), 5799 (+Cap+0nM eIF4E), 4296 (+Cap+200nM eIF4E), and 10790 (+Cap+500nM eIF4E). (*C, D*) Cumulative probability distribution (*C*) and mean values (*D*) of Nmax. Mean ±S.E.: 1.94 ± 0.02 (-Cap+500nM eIF4E), 2.05 ± 0.02 (+Cap+0nM eIF4E), 2.21 ± 0.03(+Cap+200nM eIF4E), 2.66 ± 0.02 (+Cap+500nM eIF4E). Numbers of trajectories analyzed are the same as in (*A, B*). (*E, F*) Cumulative probability distribution (*E*) and mean values (*F*) of T_1_. Mean ± S.E. (sec): 3300 ± 20 (-Cap+500nM eIF4E), 3280 ± 20 (+Cap+0nM eIF4E), 3200 ± 30(+Cap+200nM eIF4E), 2740 ± 20 (+Cap +500nM eIF4E). Numbers of trajectories analyzed are the same as in (*A, B*). (*G, H*) Cumulative probability distribution (*G*) and mean values (*H*) of ΔT_n_. Mean ± S.E. (sec): 280 ± 4 (-Cap+500nM eIF4E), 294 ± 4 (+Cap+0nM eIF4E), 240 ± 3(+Cap+200nM eIF4E), 232 ± 2 (+Cap +500nM eIF4E). Numbers of events analyzed: n = 19637 (-Cap+500nM eIF4E), 13809 (+Cap+0nM eIF4E), 12925 (+Cap+200nM eIF4E), and 49012(+Cap +500nM eIF4E). (*I, J*) Cumulative probability distribution (*I*) and mean values (*J*) of D-active. Mean ± S.E. (sec): 1260 ± 20 (-Cap+500nM eIF4E), 1200 ± 20 (+Cap+0nM eIF4E), 1160± 20 (+Cap+200nM eIF4E), 1091 ± 9 (+Cap +500nM eIF4E). Numbers of events analyzed: n = 6183 (-Cap+500nM eIF4E), 5799 (+Cap+0nM eIF4E), 6172 (+Cap+200nM eIF4E), and 21224 (+Cap +500nM eIF4E).

Compared to the Cy3-αFLAG first arrival time without eIF4E supplementation, first arrival times for the translation of the capped mRNA were decreased by 2.4% and 16.3% in extract supplemented with 200 nM and 500 nM eIF4E, respectively (Figures 3E and 3F). The Cy3-αFLAG first arrival times were similar with capped *m*^*7*^*G-3xFLAG-Fluc-pA* mRNA translation without eIF4E supplementation and with uncapped ppp*-3xFLAG-Fluc-pA* mRNA translation with 500 nM eIF4E supplementation. These results indicate that in this HeLa extract, m^7^G cap-eIF4E interactions enhanced the first-round initiation efficiency of *m*^*7*^*G-3xFLAG-Fluc-pA* mRNA translation. ΔTn and D-active analysis show that the Cy3-αFLAG binding time lags and the duration of mRNA engaged in polysome formation were similar without and with eIF4E supplementation and for both capped and uncapped mRNAs (Figures 3G-J). Collectively these results indicate that the mechanism of the regulation of initiation and polysome formation by m^7^G cap-eIF4E interactions is similar in both WGE and HeLa extract translation of *m*^*7*^*G-3xFLAG-Fluc-pA* mRNA.

## Discussion

The roles of the m^7^G cap and poly(A) tail have been previously investigated for effects on initiation but not for effects on polysome formation kinetics (23-25). By applying a single-molecule approach to track individual ribosomes during active translation, we examined effects of m^7^G cap-eIF4E and PABP-poly(A) tail interactions on polysome formation kinetics. Our results show that the m^7^G cap and eIF4E enhanced total translation activity, polysome formation, first round initiation kinetics, and the duration of active polysome formation in wheat germ and HeLa extracts. While our results confirm previous studies indicating that m^7^G cap-eIF4E interactions enhance translation (25-27), we did not observe significant m^7^G cap or eIF4E effects on initiation rates that occurred subsequent to the first-round initiation event. Our results indicate that m^7^G cap-eIF4E interactions specifically stimulate the first-round of initiation and provide insights into the role of eIF4F interactions on translation dynamics.

Interactions involving the eIF4F factors, eIF4E, eIF4A, and eIF4G, are thought to enhance translation efficiency, at least in part, by promoting a closed loop mRNA conformation (4, 5, 26). m^7^G cap-eIF4E interactions are one of several redundant eIF4F interactions that contribute to the mRNA closed loop structure and eIF4G alone is involved in multiple interactions that contribute to a closed loop conformation (4). Although eIF4G can interact with multiple partners, such as eIF4E, eIF4A, PABP, and mRNA, not all of these interactions appear necessary for mRNA to form a closed loop. Previous studies suggest that eIF4G-PABP interactions may compensate for compromised eIF4G-eIF4E interactions (28) and eIF4G-mRNA interactions may be sufficient for closed loop formation (4). Thus, independent of m^7^G cap-eIF4E interactions, closed loop formation and efficient initiation may occur through redundant eIF4G interactions. These interactions may provide an explanation for our single-molecule observations of efficient m^7^G cap-independent post first-round initiation rates. Our results suggest that, even in the absence m^7^G cap-eIF4E interactions, first-round initiation events can prime mRNA, possibly by establishing eIF4G-mRNA interactions, to maintain translationally active closed loop mRNA structures.

The poly(A) tail enhanced total translation activity, polysome formation, first round initiation kinetics, and the duration of active polysome formation (Figures 3A-F, 3I, and 3J). The observation that the m^7^G cap was required for these poly(A) tail effects supports previous studies indicating that poly(A) tail-PABP and m^7^G cap -eIF4E interactions cooperate to enhance translation (25-27). Although poly(A) tail-PABP interactions can connect mRNA 5’ and 3’ ends to form a closed loop through poly(A) tail-PABP-eIF4F-m^7^G cap interactions (4, 5), PABP may connect eIF4F-m^7^G cap interactions to mRNA sequences that are not polyadenylated. Previous studies showed that PABP can bind to unadenylated mRNA sequences (29), which could enable closed loop formation via 3’ mRNA-PABP-eIF4F-m^7^G cap interactions. In addition, closed loop formation could occur independent of a poly(A) tail by direct eIF4G binding to mRNA 3’ sequences to form 3’ mRNA-eIF4G-eIF4E-m^7^G cap interactions (4). Our results support the notion that RNA binding motifs in the N-terminus, C-terminus, and central regions of eIF4G enable redundant eIF4G-mRNA interactions that contribute to closed loop formation of mRNAs without an m^7^G cap and/or poly(A) tail. Such eIF4G interactions may contribute to previous cryo-electron tomography observations of circularized polysomes on uncapped and unadenylated mRNA (12). Our findings indicate that, on polysome forming mRNAs, the m^7^G cap and poly(A) tail serve critical roles for stimulating first-round initiation events and suggest a mechanism wherein first-round initiation activates mRNAs for efficient subsequent rounds of polysome-forming initiation events.

## Materials and Methods

### RNA synthesis

The plasmid encoding *3xFLAG-Fluc* mRNAs was as described before(30). The sequence of hairpin insert CGGCCCGCCGGCCGATATCACGGCCGGCGGGCCG was ligated to the *3xFLAG-Fluc* plasmid backbone at the 90 nt position from the 5’ end of the RNA sequence to generate *m*^*7*^*G-hp-3xFLAG-Fluc-pA* mRNA. The *3xFLAG-Fluc* plasmid was linearized with EcoRI for *in vitro* transcription of mRNAs with poly(A) tails. DNA templates for transcription of mRNAs without poly(A) tails were generated by *3xFLAG-Fluc* plasmid digestion with XbaI. Truncated construct *m*^*7*^*G-3xFLAG-truncated* was prepared from a PCR reaction with the *3xFLAG-Fluc* plasmid as a template, Forward primer 5’-CAAGGAATGGTGCATGCTAA-3’, and reverse primer 5’-CTCAGCGTAAGTGATGTCCAC-3’. RNAs were synthesized with the MEGAscript™ T7 Transcription Kit (AM1334). 5’-end capping and 3’-end biotinylation were performed using the Vaccinia Capping System (New England Biolabs) and the Pierce RNA 3’ End Biotinylation Kit (Thermo Scientific), respectively. All mRNAs were phenol-chloroform extracted and subsequently purified with the Direct-Zol RNA kit (Zymo Research), from which mRNA was eluted in water. Integrity and concentration of mRNA were assessed using denaturing (8M urea) 5% acrylamide gel electrophoresis and SYBR green II RNA staining.

### Preparation of ribosome-depleted wheat germ extract

Ribosome depleted wheat germ extract (RdWGE) was generated by centrifugation of wheat germ extract (Promega) for two rounds, each at 90,000 rpm for 1.5 hour at 4 °C with a TLA100.2 rotor and collection of the supernatant(31). WGE with partially depleted ribosomes was obtained by mixing the starting WGE and RdWGE.

### Chemicals and cap analogue

Harringtonine (vendor: LKT, catalog No.: ab141941) was stored at -20 °C with a stock concentration of 100mM in DMSO. The final concentration of harringtonine in translation reactions was 0.5 uM. The DMSO amount in final reactions was 1% of the total reaction volume. The Cy3 labeled anti-FLAG antibody (Cy3-αFLAG) concentration in translation reactions was 134 nM. Puromycin was purchased from Sigma (P7255). The final puromycin concentration used in translation reactions was 1 mM.

### The assembly of the wheat germ extract translation mixtures for single-molecule detection

Wheat germ extract with micrococcal nuclease treatment was purchased from Promega (L4380). The translation reaction mixture for single-molecule detection was assembled according to the manufacturer’s instructions with the following modifications: 1) the RNase inhibitor RNaseIn (Promega) was used at a final concentration of 0.8 U/ul; 2) the KOAc concentration was adjusted to 103 mM for optimal translation activity; 3) no mRNA was added to the reaction directly; mRNAs were 3’ end-tethered to the detection surface; 4) Cy3-αFLAG (Sigma A9594; stored at -20 °C in 50% glycerol) was added to a final concentration of 134 nM; and 5) extract was aliquotted for single-use, flash freezed, and stored at -80 °C.

### Single molecule imaging and data analysis

Construction of the single-molecule detection chamber was as described before(32). The flow channel was first incubated with 0.2 mg/ml streptavidin (Thermo scientific) for 10 min and then washed twice, each with 15 ul T50 buffer (20 mM Tris HCl, pH 7.0, 50 mM NaCl). The 3’end biotinylated reporter mRNAs in T50 buffer were added to the flow chambers, incubated for 12 min, and washed with translation buffer (20mM HEPES-KOH, 100 mM KOAc, 2.1 mM Mg(OAc)_2_) to remove unbound mRNAs. The concentration of the reporter mRNA (typically 0.25-1.3 ng/uL) was adjusted to allow single-molecule density of antibody binding. A few seconds after data acquisition, 20 uL of translation mixture was delivered through a home-built microfluidic adaptor into the channel by a Harvard Apparatus syringe pump at a speed of 150 uL/min. Translation mixture was pre-heated at 26 °C for 10 min before use. Single-molecule experiments were carried out at 29 °C in a temperature-maintaining incubator. After single-molecule imaging was completed, the translation mixture was pipetted out of the flow channel and supplemented with luciferase activity assay reagents for luminescence measurement.

Data were recorded as a kinetic series at the speed of 1 second per frame. Image analysis was performed with custom written Python and Matlab codes. Each data movie was drift and background corrected. The bound antibodies in each frame were identified based on their fluorescence intensities. The intensity signal was filtered with non-linear filtering (33). Bayesian change point detection was used with a log-normal distribution to identify transition points on single-molecule trajectories (34).

## ACKNOWLEDGMENTS

We thank Samie R Jaffrey and Ben R Hawley for providing the human eIF4E protein. This work was supported by the National Institutes of Health [R01GM121847], the Memorial Sloan Kettering Cancer Center (MSKCC) Support Grant/Core Grant [P30 CA008748], and the MSKCC Functional Genomics Initiative.

