## Supplementary Material for "m^7^G cap-eIF4E interaction stimulates polysome formation by enhancing first-round initiation kinetics"

**Figure S1. Survival probability of pre-existing Cy3-αFLAG/peptide/translating ribosome complexes during WGE translation with normal condition (black) and an addition of 1mM puromycin (dashed).**

The probability distribution was fit to a single- exponential distribution (red curves) $f(x) = a * \exp\left(-\frac{x}{t}\right) + c$, which yielded a time constant of 646 ± 6 (S.E.) sec. for the normal condition, and 54.2 ± 0.9 (S.E.) sec for the puromycin condition.

**Figure S2. Representative single molecule trajectories for WGE translation of *3xFLAG-Fluc* mRNAs**

(*A*) Raw and digitized data are shown as in gray and black, respectively for +Cap+100PA, +Cap-PA, -Cap+100PA, -Cap-PA, and 14bp *3xFLAG-Fluc* mRNAs (from top to bottom). (*B*) The mean number of co-existing antibodies on single mRNAs over time for different mRNA constructs. Numbers of trajectories analyzed: n = 2192 (+Cap+100PA), 2060 (+Cap-PA), 1865 (-Cap+100PA), 1381 (-Cap-PA), and 3405 (14bp).

**Figure S3. Schematic of kinetic parameter quantification of single-molecule trajectories for characterizing polysome formation kinetics.**

The early background intensity increase is due to delivery of translation mix supplemented with Cy3-αFLAG. When surface-anchored reporter mRNA gets activated and engaged in translation, the fluorescent intensity increases instantaneously when nascent peptides is bound by Cy3-αFLAG (green arrow) and decreases instantaneously when the Cy3-αFLAG/peptide is released from mRNA upon translation termination (blue arrow). The order of antibody binding/termination is noted on the trajectory. The definition of kinetic parameters $T_1$, $\Delta T_n$ and D-active are illustrated on the trajectory, while the definition of $N_{max}$ and $N_{total}$ are explained under the trajectory.

### Figure S1

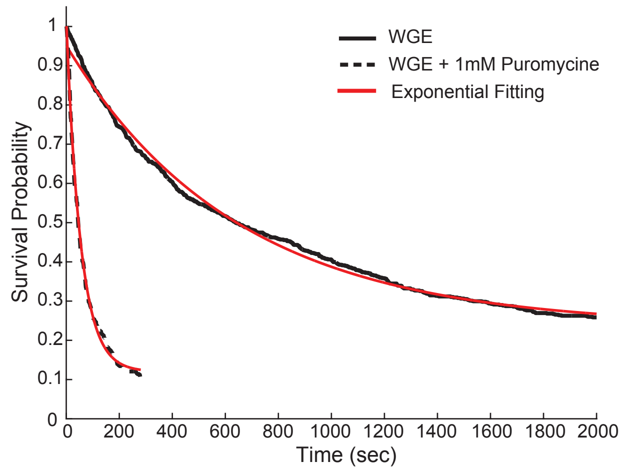
WGE
WGE + 1mM Puromycine
Exponential Fitting
Survival Probability
Time (sec)
0
0.1
0.2
0.3
0.4
0.5
0.6
0.7
0.8
0.9
1
0
200
400
600
800
1000
1200
1400
1600
1800
2000

### Figure S2

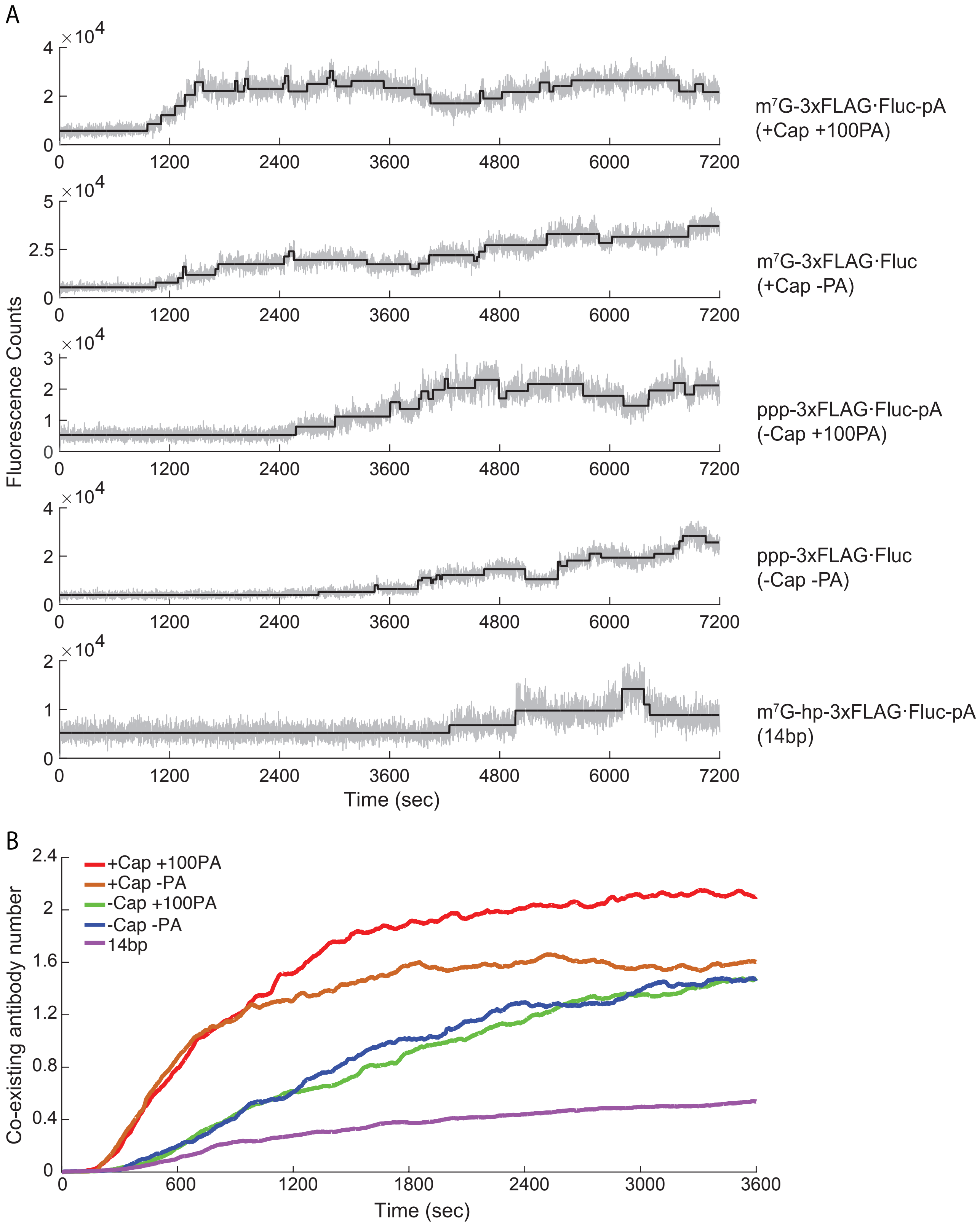

A
B
Fluorescence Counts
×10^4
m7G-3xFLAG·Fluc-pA
(+Cap +100PA)
m7G-3xFLAG·Fluc
(+Cap -PA)
ppp-3xFLAG·Fluc-pA
(-Cap +100PA)
ppp-3xFLAG·Fluc
(-Cap -PA)
m7G-hp-3xFLAG·Fluc-pA
(14bp)
Time (sec)
Co-existing antibody number
+Cap +100PA
+Cap -PA
-Cap +100PA
-Cap -PA
14bp
Time (sec)

### Figure S3

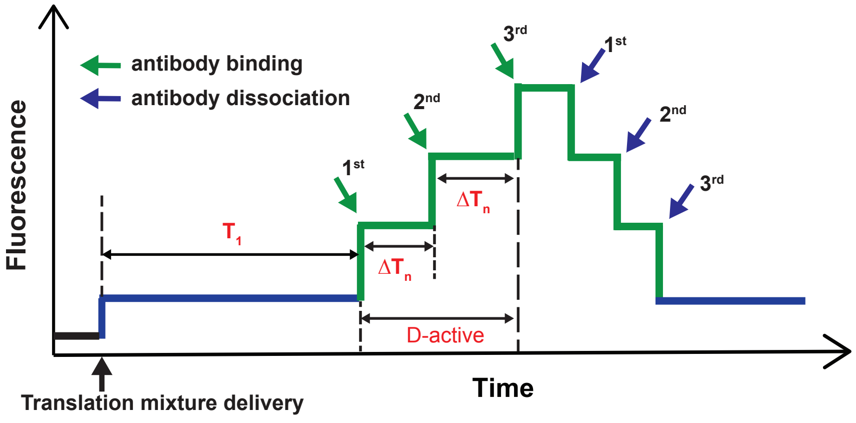


$N_{total}$ : Total number of antibodies recruited on single mRNA

$N_{max}$ : Maximum number of co-existing antibodies on single mRNA

D-active : Consecutive antibody binding duration
